# Delivering on Biden’s 2030 conservation commitment

**DOI:** 10.1101/2021.02.28.433244

**Authors:** B. Alexander Simmons, Christoph Nolte, Jennifer McGowan

## Abstract

On January 27, 2021, President Biden signed an executive order, *Tackling the Climate Crisis at Home and Abroad*, committing the United States to various goals within his campaign’s major climate policy, the *Biden Plan for a Clean Energy Revolution and Environmental Justice*. Included in this executive order is a commitment to “conserving at least 30 percent of [the United States’] lands and oceans by 2030.” This ambitious conservation target signals a promising direction for biodiversity in the United States. However, while the executive order outlines several goals for climate mitigation, the ‘30×30’ target remains vague in its objectives, actions, and implementation strategies for protecting biodiversity. Biodiversity urgently needs effective conservation action, but it remains unclear *where* and *what* this 30% target will be applied to. Achieving different climate and biodiversity objectives will require different strategies and, in combination with the associated costs of implementation, will lead to different priority areas for conservation actions. Here, we illustrate what the 30% target could look like across four objectives reflective of the ambitious goals outlined in the executive order. We compile several variations of terrestrial protected area networks guided by these different objectives and examine the trade-offs in costs, ecosystem representation, and climate mitigation potential between each. We find little congruence in priority areas across objectives, emphasizing just how crucial it will be for the Biden administration to develop clear objectives and establish appropriate performance metrics from the outset to maximize both conservation and climate outcomes in support of the 30×30 target. We discuss important considerations that must guide the administration’s conservation strategies in order to ensure meaningful conservation outcomes can be achieved over the next decade.

## Introduction

President Joseph R. Biden, Jr. has promised to usher the United States into a new era of national environmental sustainability. In his latest executive order, *Tackling the Climate Crisis at Home and Abroad*, signed on January 27, 2021, the administration will “advance conservation, agriculture, and reforestation” by committing to the goal of “conserving at least 30 percent of our lands and oceans by 2030” (EOP 2021). Furthermore, the executive order establishes the Civilian Climate Corps Initiative, which will facilitate this goal by generating new job opportunities focused on “conserving and restoring public lands and waters, increasing reforestation, increasing carbon sequestration in the agricultural sector, protecting biodiversity, improving access to recreation, and addressing the changing climate” (EOP 2021).

This target aligns with recent global commitments to protect 30% of the world’s terrestrial and marine ecosystems as part of the 2030 Agenda for Sustainable Development, known as the ‘30×30’ goal (WWF 2020). Many components of the executive order are explicit in their goals; however, the target for biodiversity conservation remains vague in its objectives, actions, and implementation strategies. Biodiversity urgently needs effective conservation action, but expectations of *where* and *what* this 30% target applies to remain uncertain amidst simultaneous—and potentially competing—goals for climate mitigation.

To address this, we encourage a systematic conservation planning framework be adopted early to ensure the 30×30 goal will achieve meaningful conservation outcomes. Such a framework will support the Biden administration’s target by enabling an inclusive process to develop explicit, quantifiable biodiversity and climate objectives that will guide the placement of conservation strategies where they benefit nature most, and minimize negative impacts on people, communities, and industries. Using this framework, the incoming administration is presented with an exceptional opportunity to develop a transparent, systematic, science-based, and community-informed framework to deliver on national conservation commitments and pioneer a global standard for achieving the 30×30 goal.

### Protected areas and the biodiversity crisis

What is considered ‘protected’ in the US is subject to interpretation. According to international reporting standards of the United Nations Environment Programme (UNEP), terrestrial protected areas currently cover nearly 12% (1.12 M km^2^) of US lands (UNEP-WCMC 2020). However, the official national inventory—the Protected Area Database of the United States (PAD-US)—is far more inclusive of what is considered ‘protected.’ The most recent PAD-US data considers more than 31% of land under various forms of protection, including 13% (1.25 M km^2^) with strict mandates for biodiversity protection (PAD GAP status 1 and 2), and an additional 18% (1.67 M km^2^) protected from conversion yet subject to multiple permissible uses (PAD GAP status 3), such as logging and mining (USGS GAP 2020) (Fig. 1). The Biden administration must determine what baseline it will consider for achieving this 30×30 target; under the most exclusive baseline with greatest biodiversity protection, the coverage of terrestrial protected areas may need to expand more than twice its current size within the next decade—a welcomed, albeit ambitious, target.

**Fig. 1.**
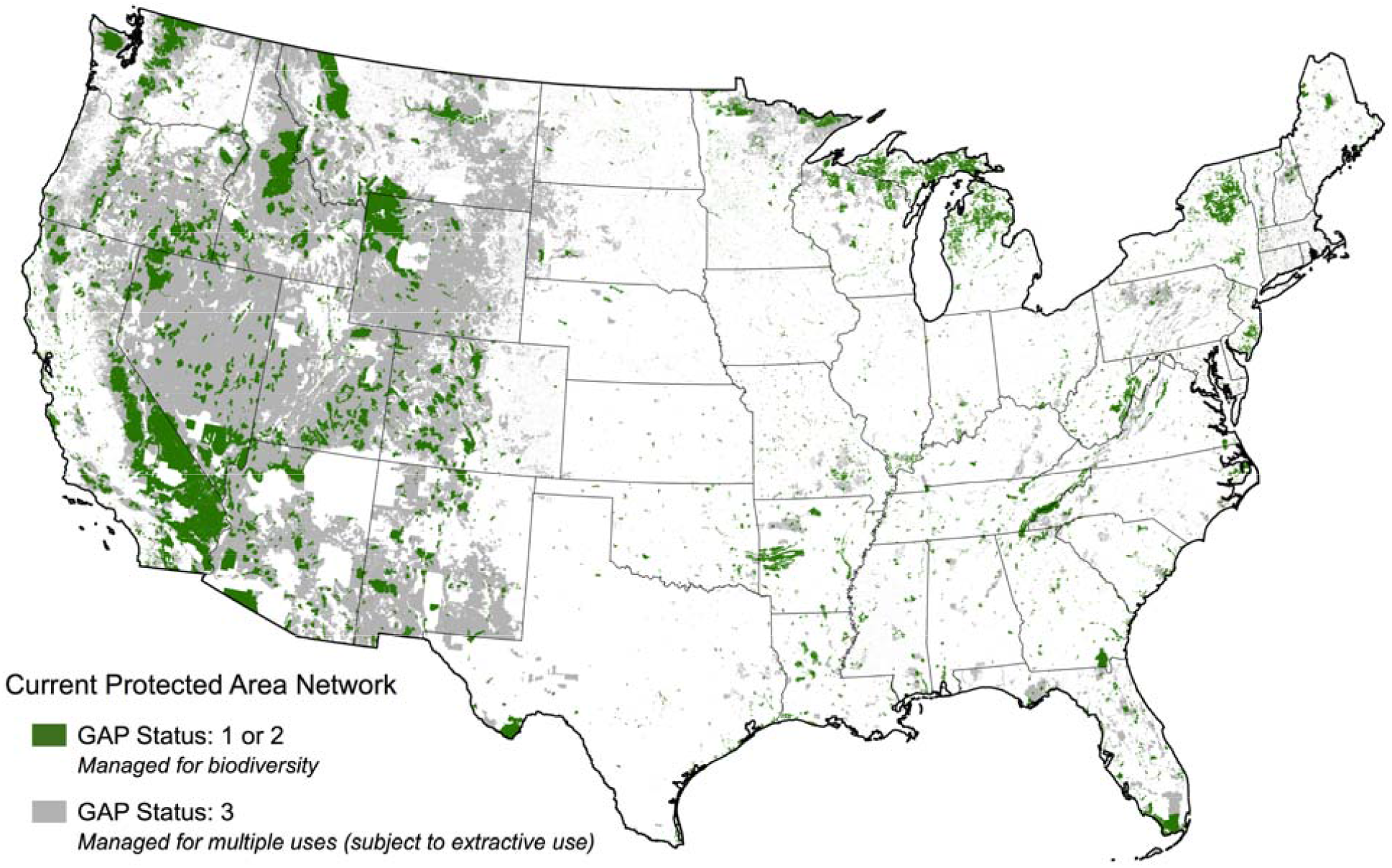
Current distribution of terrestrial protected areas with known mandates for biodiversity protection on undeveloped land in the conterminous United States. Protected areas are distinguished by Gap Analysis Project (GAP) status codes. Data obtained from the Protected Areas Database of the United States (USGS GAP 2020).

The current protected area network is insufficient to curtail significant biodiversity losses. Recent estimates suggest one-third of terrestrial species in the US are threatened with extinction, of which just 11% have adequate representation within existing protected areas (Dietz et al. 2020). There is a large bias toward protecting lands and ecosystems in Alaska and other remote, sparsely inhabited areas where competition with agriculture is low (Bargelt et al. 2020; Venter et al. 2017). The concentration of protected areas in the western conterminous US contrasts the distribution of endemic species in the southeast (Jenkins et al. 2015), where protected areas are few in number and small in size (Venter et al. 2017).

Furthermore, the future of protected areas in the US is increasingly uncertain. Protected area downgrading, downsizing, and degazettement (PADDD) has impacted more than 0.5 M km^2^ of protected lands in the US, with almost an equivalent 0.4 M km^2^ of additional land threatened by PADDD proposals brought forth in the last 20 years alone (Kroner et al. 2019); most notably, the reductions of Bears Ears (85%) and Grand Staircase-Escalante National Monuments (51%) in 2017 under the Trump administration constitute the largest downsizing events in US history (Kroner et al. 2019). Even if existing protected areas could be secured into the future, it is likely that climate change will jeopardize the effectiveness of these lands for biodiversity without adaptive and proactive management. Due to their geographic bias, existing national parks are more vulnerable to climate change than unprotected lands in the US (Gonzalez et al. 2018). Areas with greater potential to serve as species- and climate-refugia in the future offer exceptional conservation value, yet many of these important areas are currently unprotected (Lawler et al. 2020; Stralberg et al. 2020).

### One target, multiple potential objectives

Without explicit objectives, it is unclear how the 30×30 target will achieve Biden’s goals of biodiversity protection and climate mitigation. As observed in the global response to the Convention on Biological Diversity’s previous Aichi Target 11 (protection of 17% terrestrial and 10% marine ecosystems globally), area-based protection targets are susceptible to inadequate and inequitable placement, underachievement, insufficient resourcing, and other perverse outcomes as countries aim to quickly and cheaply increase the quantity of ‘protected’ lands and waters (Barnes et al. 2018). Achieving different objectives will require different conservation strategies and, in combination with the associated costs of implementation, will lead to different priority areas for conservation actions. The most affordable locations may not provide the most climate mitigation potential, and areas with the most climate mitigation potential may not adequately secure threatened species from extinction. Without systematic planning, the potential for synergies between objectives may not be fully realized, jeopardizing efficiency and missing critical opportunities to provide evidence that biodiversity and climate goals can be equitably achieved alongside sustainable management and economic growth on land and sea.

To illustrate the importance of early, definitive objective-setting for the Biden administration’s forthcoming conservation planning, we show how meeting different objectives will drive priorities towards disparate geographies within the US, delivering variable outcomes for biodiversity and climate goals. We identified cost-effective expansions of the existing protected area network to fully protect 30% of undeveloped land under four objectives reflective of the goals in the executive order: (1) area-based objective, (2) landscape-based objective, (3) species-based objective, and (4) carbon-based objective. While we acknowledge the 30×30 goal will be met through a combination of land, freshwater, and marine conservation, we focused this illustrative example on meeting the 30% target within the conterminous US landscape where we have the best available ecological and land value data. Understanding and quantifying requisite trade-offs will be critical to this administration’s conservation decision-making and will require identifying relevant performance metrics in tandem with objective setting. To highlight this, we compare the performance of each objective according to three network-level performance metrics: total cost, ecosystem representation, and climate mitigation potential.

## Methods

We divided the conterminous US into the same 100 km^2^ planning units as Lawler et al. (2020), for a total of 79,784 planning units covering all terrestrial areas. We excluded developed areas from potential selection and from our estimates of the area available to reach the 30% target. These developed areas include all land classified by the 2016 National Land Cover Database as ‘developed, open space’, ‘developed, low intensity’, ‘developed, medium intensity’, and ‘developed, high intensity’ (Yang et al. 2018). We further excluded all undeveloped land classified as a protected area under GAP 1 or 2 protection status (USGS GAP 2020) from potential selection. We do not exclude undeveloped land classified under GAP 3 protection status for the following reasons: (1) these protected areas increase the existing protected area coverage above 30% of the U.S. (Fig. 1), so they (or at least some) are unlikely to be considered in the baseline by the Biden administration, (2) they do not have such strict biodiversity protection mandates as GAP 1 and 2 protected areas, and (3) the permissible uses (e.g. logging and mining) introduce large variation in the potential impacts on biodiversity between GAP 3 protected areas.

Approximately 574,412 km^2^ (7.49%) of the conterminous U.S. is protected under GAP 1 and 2; therefore, we required at least 1,723,452 km^2^ (22.51%) of undeveloped land to be selected for each objective in order to reach the 30% target. Per common practice in systematic conservation planning, all planning units with more than 50% of their total area classified as a GAP 1 or 2 protected area were excluded from potential selection, including any remaining unprotected and undeveloped land within the respective planning units. For our illustrative purposes, we cost-effectively selected the additional 22.5% of lands for each objective based upon the most conservative assumption of full protection through land acquisitions without residual extractive uses, such as timber or grazing. We used the most recent high-resolution estimates of the 2010 fair market value of private lands in the conterminous U.S. (Nolte 2020) to calculate the costs per hectare of undeveloped land within each planning unit. While we do not advocate for meeting the 30% target exclusively through strict protection, we use this approach to be illustrative of the upper bounds of socio-economic costs. This approach overestimates the cost of a diversified protection strategy that involves partial protection (e.g. through easements or “working” lands), yet it is likely to reflect much of the spatial heterogeneity in costs for such alternative strategies.

### Protected area expansions

For the area-based objective, we sorted all planning units available for selection according to the cost per hectare of undeveloped land within them. We progressively selected all undeveloped and unprotected lands within the planning units with the lowest cost per hectare until their cumulative area exceeded 1,723,452 km^2^. For the landscape objective, we identified all undeveloped and unprotected land overlapping with the Resilient and Connected Network (RCN) of landscapes produced by The Nature Conservancy (TNC 2018). We included lands classified under all combinations of the RCN—’resilience and flow’, ‘resilience and recognized biodiversity’, and ‘resilience, flow, and recognized biodiversity’—which cover 2,158,031 km^2^ (28.19%) of undeveloped and unprotected land considered in this analysis. Areas classified as tribal lands were not available for inclusion in the RCN data. We followed a consistent approach as the area-based objective for selecting new protected areas: we limited the selection opportunities to all planning units containing undeveloped and unprotected land classified within the RCN, and progressively selected areas with the lowest cost per hectare until meeting the cumulative area target.

For the species-based objective, we use methods and species data from Lawler et al. (2020) to identify cost-effective protected area networks for species conservation under climate change. The conservation prioritization is formulated as a *minimum set* problem – which identifies the set of planning units that most cost-effectively achieves a predefined set of species-specific targets – and solved it with the Marxan conservation planning software (Ball et al. 2009). We base our analysis on the most comprehensive scenario of the original study (“all”), which includes protection targets for 1,460 current and future species distributions, 100% of climatic refugia, and 20% of climate corridors. In line with the analytical framework of our study, we only consider species presence on undeveloped land in each planning unit. To achieve 30% protected area coverage for the contiguous U.S., we scale species-specific protection targets as a function of species range using an inverse hyperbolic sine transformation:

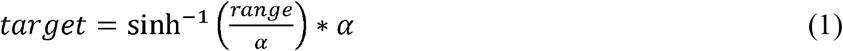

This function has similar properties as the transformation function proposed by Rodriguez et al. (2004) for global species conservation planning—namely, targets that start at 100% of range size for species with small ranges, with percentages gradually declining as species ranges increase. Here, *a* is a scaling parameter, which we adapt iteratively until the optimization returned 30 ± 1.0% coverage for the conterminous US (*α*= 21000). The final cumulative area covered 30.69% (2.35 M km^2^) of the study area, slightly higher than the 30.00% of the other objectives.

For the carbon-based objective, we prioritize protection of grasslands and forest at risk of being converted to another land use. We obtained high-resolution maps of remnant forests and grasslands and shrublands in the conterminous US from Fargione et al. (2018). In their study, Fargione et al. (2018) estimated future forest and grassland/shrubland conversion risk based upon conversion rates of different types of vegetation during 1986-2000 (forest vegetation) and 2008-2012 (grassland/shrubland vegetation). Conversion rates are based upon vegetation clearance resulting in a change in land use; this does not include vegetation clearing where the land use does not change (e.g. forest clearance as part of timber rotations). All grasslands were considered at-risk of conversion, but due to the low rates of past forest conversion, only the top 25% of forest vegetation types converted in the past were considered at high risk of conversion in the near future—see Fargione et al. (2018) for details on the methodology. We overlapped these maps with undeveloped and unprotected lands used in this study to identify areas available for protection within grasslands/shrublands, high-risk forests, and all other (low-risk) forests. All planning units containing undeveloped and unprotected grassland/shrubland or high-risk forest were selected for protection regardless of costs. In total, these areas accounted for 387,333 km^2^ (5.06%) of all undeveloped and unprotected land, costing $458 billion ($11,816 ha^-1^). To reach the 30% target at minimum cost, we then progressively selected areas containing low-risk forest with the lowest cost per hectare until meeting the cumulative area target.

### Performance metrics

To compare potential costs, we calculated the total sum of the costs of undeveloped land selected for each objective based on the 2010 fair market value data (Nolte 2020) used to select the cheapest undeveloped private lands for each objective, as described previously. To calculate ecosystem representation within the new protected area network of each objective, we obtained the most recent map of world ecosystems (Sayre et al. 2020) and excluded all ecosystems classified as ‘converted’ from their natural state. A total of 148 ‘natural’ ecosystems were included in the analysis. We overlapped these natural ecosystems with all undeveloped and unprotected land selected within each objective, as well as all land classified as GAP 1 or 2 protected areas. Areas overlapping with ‘converted’ ecosystems were not included in the representation analysis, leaving 85.73% of the area-based network, 94.05% of landscape-based network, 85.78% of species-based network, 89.59% of the carbon-based network, and 95.69% of the existing protected area network (GAP 1 or 2) available to assess ecosystem representation. To calculate the Representation Achievement Score we used the R-package “ConsTarget” (Jantke et al. 2019) which calculates the mean proportional target achievement for all biodiversity features of interest found in a conservation network or protected area estate. We calculated the score against targets of 30% for all 148 natural ecosystems using the selected area for each objective as well as the existing baseline PA network.

To estimate climate mitigation potential for each objective, we calculated the total estimated carbon emissions attributed to grasslands/shrublands and high-risk forests based upon data from Fargione et al. (2018). This spatial data estimates the per hectare carbon emissions (Mg C ha^-1^) from grasslands and shrublands, and albedo-adjusted per hectare carbon emissions (Mg C ha^-1^) for the top 25% of forests at greatest risk of conversion—see Fargione et al. (2018) for details on the methodology. We resampled the existing datasets to align with our 900 m^2^ pixels of undeveloped and unprotected land. For the grassland/shrubland dataset, we multiplied the original values (in Mg C ha^-1^) by 0.09 ha to obtain Mg C estimates per pixel (900 m^2^). For the forest dataset, we divided the original values (in dag C ha^-1^) by 100,000 and multiplied by 0.09 ha to obtain the same Mg C estimates per pixel. Emissions estimates were attributed to all undeveloped and unprotected land selected within each objective and summed to achieve the total climate mitigation potential for each objective in avoided emissions from future grassland, shrubland, and forest conversion (Gt C).

## Results

A purely area-based objective would lead to a large protection bias in the western plains and northern Great Basin, with minimal representation in the Southeast (Fig. 2a). This approach would do little to improve the existing distributional biases of the current protected area network, falling below the acceptable threshold for ecosystem representation. This objective also offers the lowest climate mitigation opportunity, potentially avoiding just 0.08 Gt C in emissions from grassland and forest conversion. While this objective presents the cheapest option for the 30×30 target, costs for complete land acquisition could still reach upwards of $270 billion ($1,567 ha^-1^). Approximately 33% of the areas selected for protection under this scenario are currently under GAP 3 protection status (Fig. 3a).

**Fig. 2.**
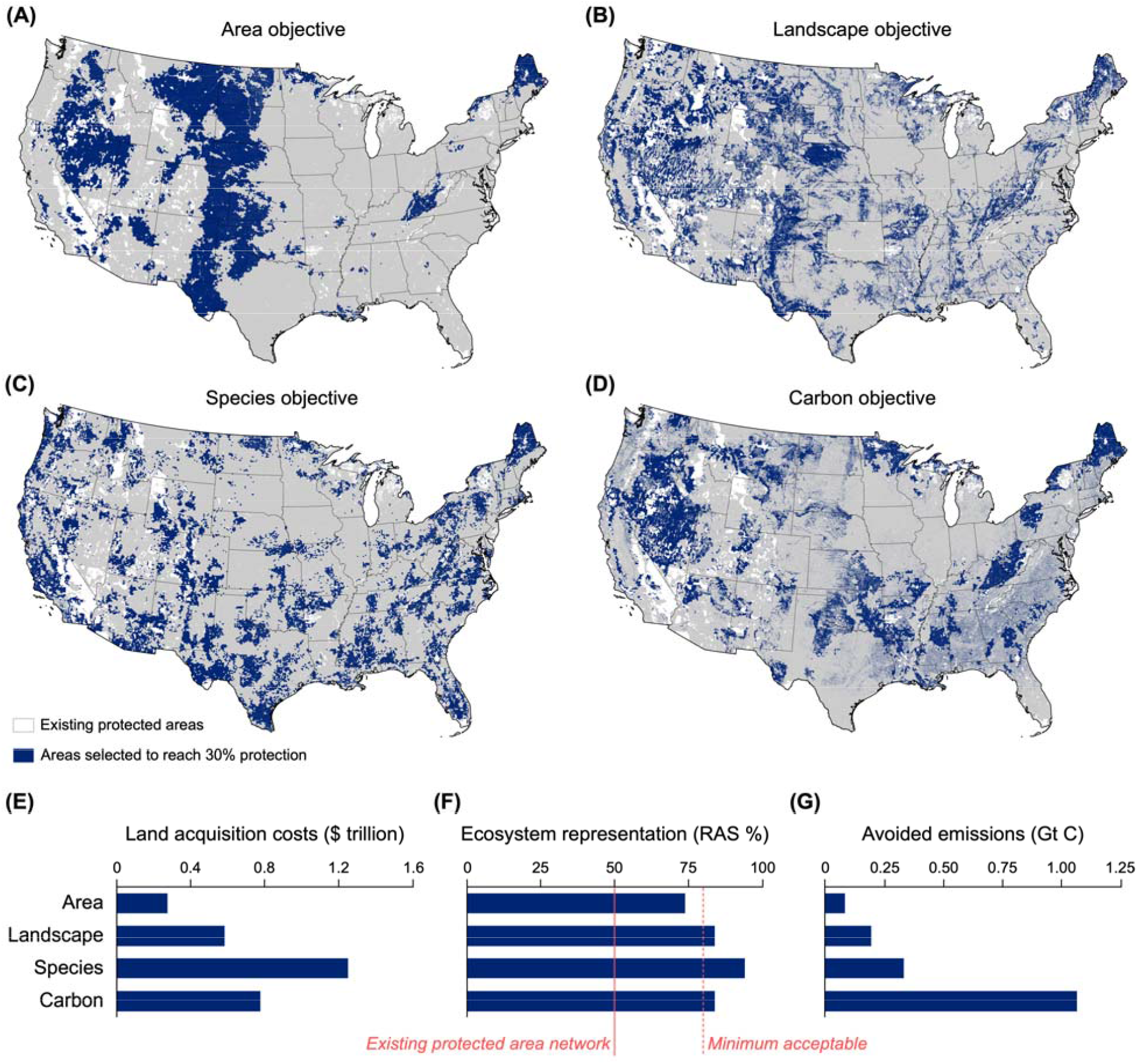
Outlook of the ‘30×30’ target under different objectives. **(A-D)** The most cost-effective areas to achieve 30% protection of land in the conterminous US according to area, landscape, species, and carbon-based objectives. **I** Total estimated land acquisition costs for areas selected in each objective. **(F)** Ecosystem representation within each objective based upon representation achievement score (RAS). **(G)** Climate mitigation potential for each objective based upon avoided emissions of grasslands, shrublands, and forests at greatest risk of future land conversion.

**Fig. 3.**
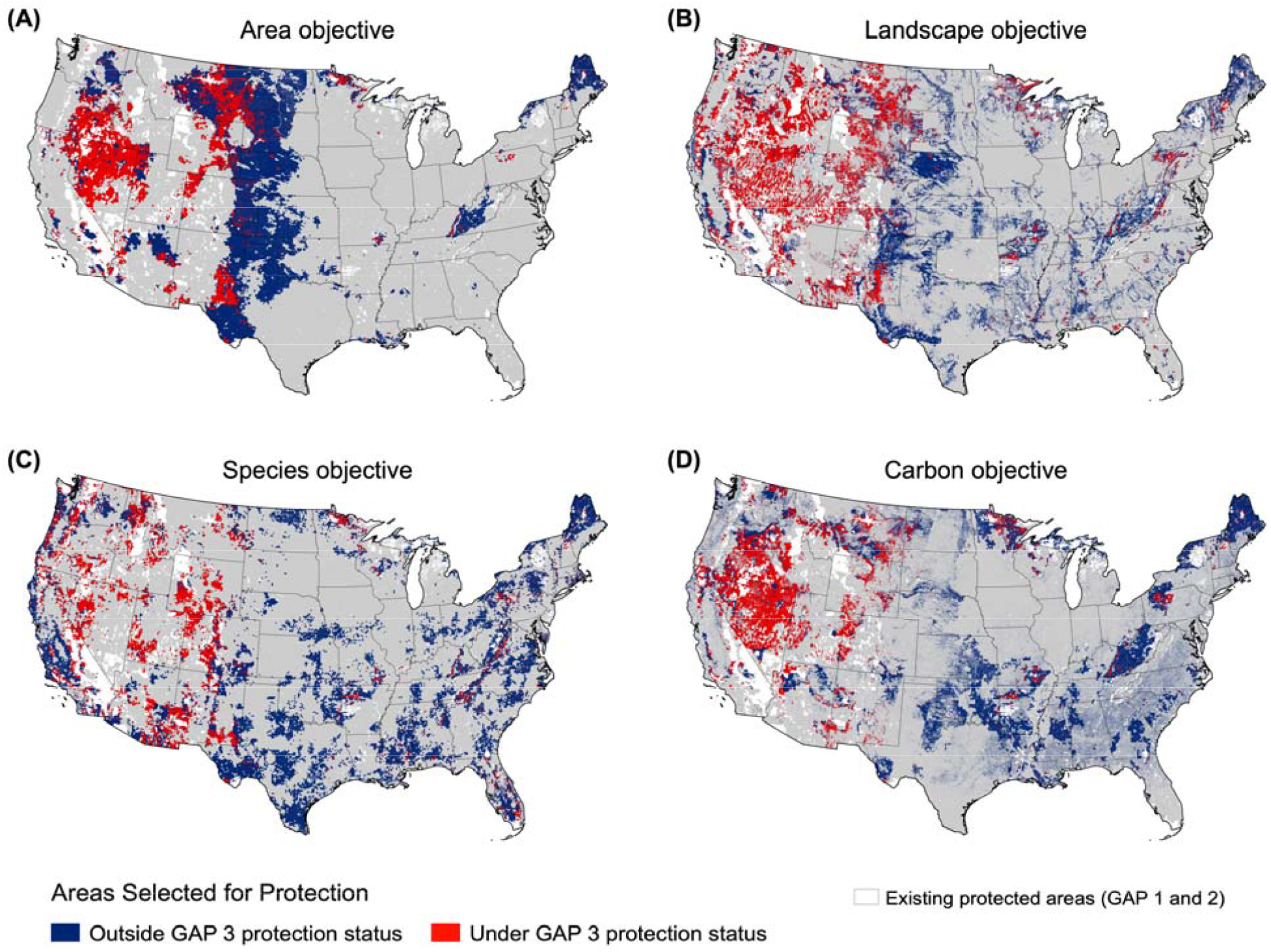
Extent of undeveloped land selected for protection across the **(a)** area, **(b)** landscape, **(c)** species, and **(d)** carbon objectives, highlighting areas currently classified as GAP 3 protected areas (red).

The landscape-based objective also has a large presence in the West, but unlike the area-based objective, it is more representative of the Southeast, and most states have some representation in the new protected area network (Fig. 2b). Though the size of individual protected areas is smaller under this objective, the network is well-connected, often consisting of long stretches of protected areas. This approach meets the minimum acceptable representation score (RAS 84%), but many of the proposed areas surround existing protected areas, where representation of these ecosystems may already be high in the baseline. The greater inclusion of grasslands and forests at risk of conversion increases the climate mitigation potential more than twice that of the area-based objective (0.20 Gt C). These ecosystem and emissions improvements, however, come at more than twice the cost of the area-based objective ($580 billion; $3,366 ha^-1^). This objective is the most inclusive of areas managed for multiple uses, with nearly 42% of selected areas currently under GAP 3 protection status (Fig. 3b).

The species-based objective produces the most representative protected area network of all objectives, with a greater presence in the eastern and southern U.S. and a smaller presence in the western plains and Great Basin where existing protected areas are concentrated (Fig. 2c). The proposed protected areas are larger but more dispersed than in the landscape objective. This objective comes closest to achieving the ecosystem representation target (RAS 94%), and this greater diversity also leads to greater climate mitigation potential (0.33 Gt C). However, these improvements come at a cost upwards of $1.25 trillion ($7,038 ha^-1^)—more than twice the cost of the landscape objective. This network is the least inclusive of areas currently under GAP 3 protection status (27%) (Fig. 3c).

Per the design of the carbon-based objective, the resulting protected area network consists of a more representative coverage of forested and grassland ecosystems, achieving an equivalent representation score as the landscape objective (RAS 84%). Because most forests threatened with conversion are within the southeast, protection is more representative of this region than all other objectives, but the higher costs of land in these areas result in smaller patches of protection across the region (Fig. 2d); elsewhere, where forests are less threatened with conversion, larger patches of protection exist on land that is exceptionally cheaper to acquire. This is the second most expensive objective ($775 billion; $4499 ha^-1^), but it would deliver the greatest climate mitigation potential (1.07 Gt C)—more than three times the species-based objective and nearly 13 times the area-based objective. Approximately 30% of these areas are under GAP 3 protection status (Fig. 3d).

Overall, we find little congruence in priority areas across objectives (Fig. 4). Areas that were selected for protection under all four objectives cover just 2% (0.15 M km^2^) of the conterminous US, primarily concentrated in the Great Basin, northern Maine, western Appalachian Plateau, and southwestern Texas. An additional 7.5% (0.57 M km^2^) of the country was selected under three objectives, and 16% (1.24 M km^2^) under two objectives. Most concerning, 28% of the country (2.16 M km^2^) was selected under just one objective, emphasizing the heterogeneity of biodiversity, ecosystems, and land-uses in the conterminous US and the challenge of finding areas that can meet diverse objectives. A large proportion of the country (39%; 2.97 M km^2^) was never selected for protection, most notably in the production-intensive Midwest and the highly developed Northeast Coast.

**Fig. 4.**
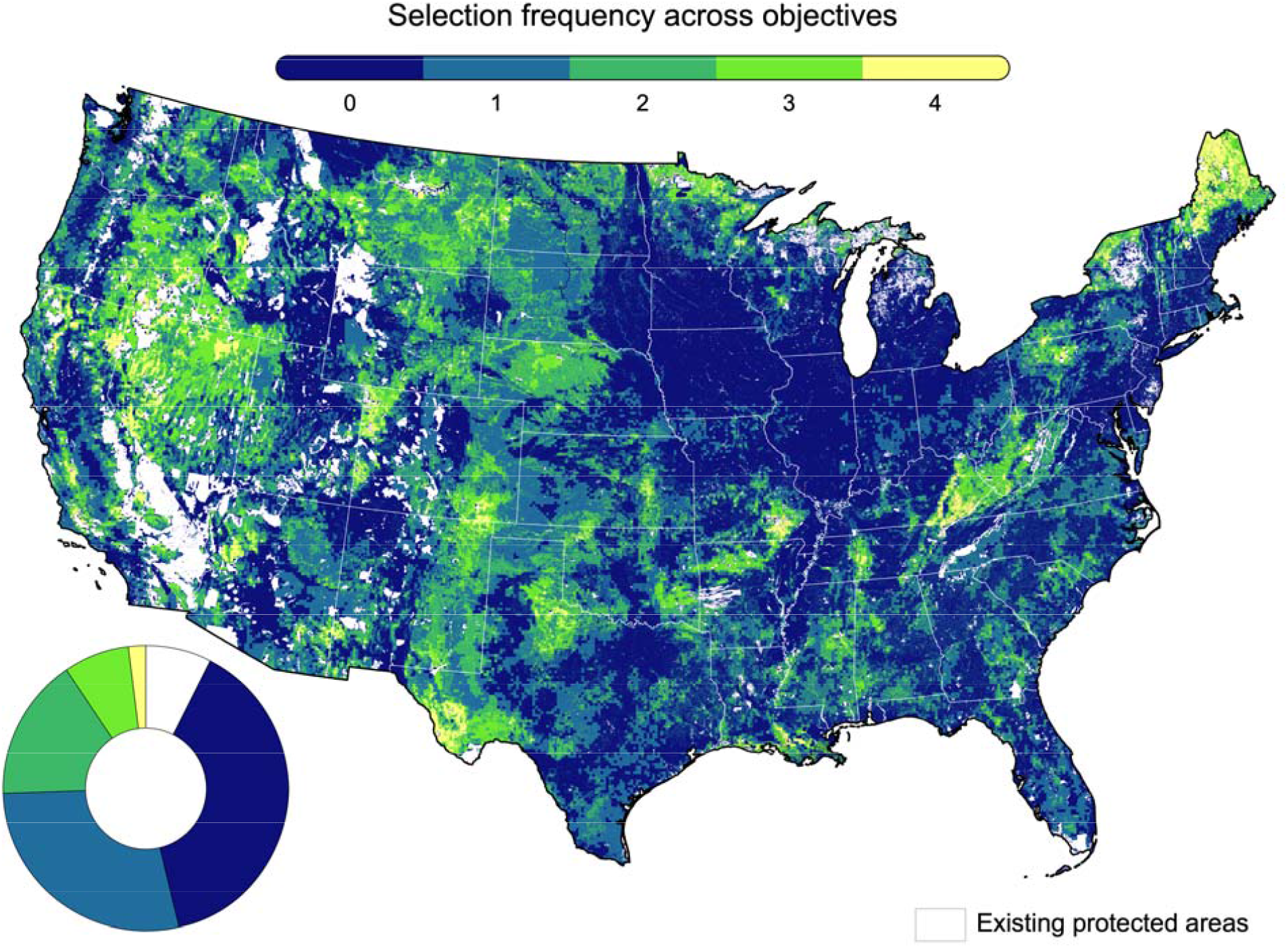
Extent and proportion of the conterminous United States selected for protection under multiple objectives.

## Discussion

The 30×30 target will not be a panacea for the United States’ conservation problems, but with the right objectives and actions, the target can be an important policy vehicle to deliver meaningful conservation and climate outcomes. Biden’s support for this international 30×30 goal is a promising signal of a return to the country’s global citizenship in the fight for conservation and climate action. While no single objective delivers the maximum benefits across all biodiversity and climate goals of the 30×30 target, the administration still has the opportunity to create positive outcomes during the next decade. However, translating this global conservation commitment into national-level actions will be challenging. We propose several considerations that will be crucial to ensuring the next decade of environmental protection is done efficiently, cost-effectively, and equitably to maximize benefits for people and nature.

### Set immediate and clear objectives to guide prioritizations of the 30% target

We have demonstrated how strategic implementation of the 30×30 target will require clear objectives to understand trade-offs and maximize conservation and climate outcomes. Yet even with the relatively simple objectives we have examined here, only 2% of the conterminous US was selected for protection under all four objectives. Contrast these limited ‘no regrets’ priorities with the 28% of lands selected for just a single objective and the trade-offs in priority areas becomes more consequential. Such a small percentage of ‘no regrets’ lands means transparency and consistency in how resource allocation decisions are made will be paramount.

It is encouraging that, with the simultaneous signing of the *Presidential Memorandum on Scientific Integrity and Evidence-Based Policymaking*, President Biden is committed to ensuring that the administrations’ decisions will be informed by “the best available science and data” (EOP 2021). Biodiversity and climate objectives for the 30×30 target will need to be guided by our best available knowledge across scientific disciplines to find solutions that can maximize benefits for species, ecosystems, landowners, industries, and our climate.

### Protect what is threatened, restore where there is opportunity

To create real impact, we must identify where the most pressing abatable threats are and where we can achieve the highest return on investment for actions that mitigate those threats (Withey et al. 2012). For example, prioritizing places with large amounts of non-threatened above-ground biomass may prove less impactful than prioritizing forests that are most likely to be converted or harvested in the coming decades. Additionally, prioritizing areas within species current distribution ranges may not generate the long-term benefits of prioritizing areas within both current and future distribution ranges under climate change. Such a strategy can facilitate the design of the 30×30 target over the next decade and avoid placing protected areas in locations under minimal threat—a characteristic that plagues the global protected area network (Joppa and Pfaff 2011).

While we have focused this outlook on protection, identifying restoration opportunities will also be important for delivering Biden’s goal of restoring public lands and waters. Similar to our present analysis, priority areas for restoration will be influenced by specific objectives, actions, costs and feasibility (Brown et al. 2015). For example, restoration in the eastern Midwest may deliver the greatest climate mitigation potential, but restoration in the Southeast and West Coast may yield the greatest benefits for biodiversity (Strassburg et al. 2020). Restoration activities can be expensive with low probabilities of success, so identifying clear strategies for resource allocation will be essential (Rohr et al. 2018). Evidence suggests that natural regeneration can lead to greater restoration success rates at lesser costs than active restoration (e.g. seeding, planting, burning) (Crouzeilles et al. 2017). Thus, the administration should consider where there are greater opportunities to achieve cost-effective and successful restoration outcomes.

### Establish appropriate performance metrics to evaluate progress and impact

Crucial to this approach will be the design of meaningful performance and evaluation protocols that can sufficiently track the progress of these interventions against their stated objectives. To date, there is no current international published guidance explicitly linked to the 30×30 agenda in this regard. Establishing a core set of meaningful indicators linked to the stated goals of the 30×30 plan from the outset will help ensure the objectives are aligned, monitored, and measured against quantifiable outcomes. Drawing from the post-2020 Biodiversity Monitoring Framework (OECD 2019) and using a broad suite of biodiversity indicators for species, ecosystems and their services, landscape connectivity, and climate would ensure that the US is aligned with international reporting obligations for biodiversity, while setting domestic precedent.

Further alignment and development of measures of social equity, inclusion, and racial and social justice will be equally critical. These considerations of “fairness” in conservation have increased over the last decade, with growing concerns over who bears the burden of conservation interventions, who is excluded from decision-making, and whose rights and interests are recognized in the process (Friedman et al. 2018). Social and culturally inclusive performance metrics should be identified that can properly evaluate impacts of protection on local communities across multiple dimensions, including economic living standards, governance and empowerment, social relations, and subjective well-being (McKinnon et al. 2016).

### Capitalize on the diversity of policy instruments for protection

Effective conservation outcomes can be achieved using many policy levers. Protected areas are just one instrument in our conservation toolkit. In the last few years, the International Union for Conservation of Nature has pushed for greater adoption of other effective area-based conservation measures (OECMs), which aim to achieve long-term biodiversity conservation under a more diverse consideration of important ecosystem services, greater recognition of local livelihoods and cultural values, and a more inclusive suite of governmental, organizational, and indigenous or community stakeholders (Laffoley et al. 2017). These bottom-up approaches to conservation recognize the contributions and knowledge of indigenous management, increase probabilities of success, inspire environmental stewardship within communities, and can be more cost-efficient to implement in the long-term.

Such mechanisms will be important to achieving the 30×30 goal for the incoming administration and should be weighed carefully against more restrictive protected areas expansion. Furthermore, collaboration between federal, state, tribal communities, NGOs, and land trusts will be required to achieve a comprehensive 30% network across the United States. The executive order’s commitment to “stakeholder engagement from agricultural and forest landowners, fishermen, Tribes, States, Territories, local officials, and others” (EOP 2021) shows that the administration is aiming for active inclusion of diverse stakeholders in implementing the target, and we hope such inclusive processes will be delivered in the coming years.

While the existing evidence base tends to favor a land-sparing approach to conservation in production landscapes (i.e. maximizing yields on existing farms and sparing surrounding lands for biodiversity) (Balmford et al. 2018), integrating conservation into “working” lands and seas will be critical for delivering positive outcomes for nature that should not be discounted in achieving the 30×30 goal. Improved management practices (e.g. longer timber rotations or improved fisheries management) have the potential to produce greater biodiversity and climate mitigation benefits (Fargione et al. 2018) for potentially less costs than establishing new protected areas. Revisiting domestic policies that subsidize harmful agriculture, fisheries and forestry activities is now recognized as one of the most impactful ways to recalibrate government expenditures to better protect biodiversity (Deutz et al. 2020).

Conservation easements, agri-environmental schemes, and other private land conservation programs have been championed globally to enhance ecosystem services in production lands and waters (Kamal et al. 2015), yet these instruments are underutilized in the United States (Bargelt et al. 2020). The executive order again shows promise that these alternative instruments will be included within the 30×30 target, with desires to increase adoption of “climate-smart agricultural practices that produce verifiable carbon reductions and sequestrations” (EOP 2021). However, the administration must also recognize the importance for biodiversity in production lands and seas, and a greater diversity of these programs should be promoted that can deliver multiple environmental benefits beyond just climate mitigation.

Finally, in some areas, significant environmental benefits could also be gained within existing protected areas. For example, 27-42% of areas selected in our different objectives are currently classified as GAP 3 protected areas (i.e. managed for multiple uses, such as logging and mining) (Fig. 3). These areas could be upgraded to GAP 1 or 2 status to offer more explicit biodiversity protection.

Delivering on Biden’s 30×30 commitment will be challenging, but several of these challenges can be mitigated using the systematic conservation planning framework we have outlined here. The executive order is a promising first step. To ensure efficient, effective, and equitable conservation outcomes can be achieved, the Biden administration must now focus on establishing clear objectives to guide prioritizations of places and actions for biodiversity protection and climate mitigation, using appropriate performance metrics to ensure interventions maximize environmental benefits and minimize perverse outcomes for people, communities, and industries. While we have focused this discussion on terrestrial systems in the United States, these issues also apply to the freshwater and ocean systems domestically and in the 84 countries that have already pledged their commitment to this global 30×30 target (WWF 2020). Countries adopting core principles of systematic conservation planning can prioritize the appropriate actions through inclusive and democratic processes to ensure cost-effective priorities are achieved within their own unique contexts. As the world watches President Biden propel the US into the next decade of climate action, we urge the administration to seize this opportunity to advance international conservation efforts and deliver smart national solutions to the escalating biodiversity and climate crises.

## Acknowledgments

We are grateful to T. Cors and K. Gallagher for providing feedback on an early version of this paper.

## Literature Cited

Ball IR, Possingham HP, Watts ME. 2009. Marxan and relatives: software for spatial conservation prioritization. Pages 185-195 in Moilanen A, Wilson KA, Possingham HP, editors. Spatial Conservation Prioritization: Quantitative Methods and Computational Tools. Oxford University Press, New York.

Balmford B, Green RE, Onial M, Phalan B, Balmford A. 2019. How imperfect can land sparing be before land sharing is more favourable for wild species? Journal of Applied Ecology 56:73–84.

Bargelt L, Fortin MJ, Murray DL. 2020. Assessing connectivity and the contribution of private lands to protected area networks in the United States. PLoS One 15:e0228946.

Barnes MD, Glew L, Wyborn C, Craigie ID. 2018. Prevent perverse outcomes from global protected area policy. Nature Ecology and Evolution 2:759–762.

Brown CJ, Bode M, Venter O, Barnes MD, McGowan J, Runge CA, Watson JEM, Possingham HP. 2015. Effective conservation requires clear objectives and prioritizing actions, not places or species. Proceedings of the National Academy of Sciences USA 112:E4342.

Crouzeilles R, Ferreira MS, Chazdon RL, Lindenmayer DB, Sansevero JBB, Monteiro L, Iribarrem A, Latawiec AE, Strassburg BBN. 2017. Ecological restoration success is higher for natural regeneration than for active restoration in tropical forests. Science Advances 3:e1701345.

Deutz A, Heal GM, Niu R, Swanson E, Townshend T, Li Z, Delmar A, Meghji A, Sethi SA, Tobin-de la Puente J. 2020. Financing nature: closing the global biodiversity financing gap. The Paulson Institute, The Nature Conservancy, and the Cornell Atkinson Center for Sustainability.

Dietz MS, Belote RT, Gage J, Hahn BA. 2020. An assessment of vulnerable wildlife, their habitats, and protected areas in the contiguous United States. Biological Conservation 248:108646.

EOP (Executive Office of the President). 2021. Executive Order 14008–Tackling the climate crisis at home and abroad. 86 FR 7619. Federal Register, Washington, DC.

Fargione JE, et al. 2018. Natural climate solutions for the United States. Science Advances 4:eaat1869.

Friedman RS, Law EA, Bennett NJ, Ives CD, Thorn JPR, Wilson KA. 2018. How just and just how? A systematic review of social equity in conservation research. Environmental Research Letters 13:053001.

Gonzalez P, Wang F, Notaro M, Vimont DJ, Williams JW. 2018. Disproportionate magnitude of climate change in the United States national parks. Environmental Research Letters 13:104001.

Jantke K, Kuempel CD, McGowan J, Chauvenet ALM, Possingham HP. 2019. Metrics for evaluating representation target achievement in protected area networks. Diversity and Distributions 25:170–174.

Jenkins CN, Van Houtan KS, Pimm SL, Sexton JO. 2015. US protected lands mismatch biodiversity priorities. Proceedings of the National Academy of Sciences USA 112:5081–5086.

Joppa LN, Pfaff A. 2011. Global protected area impacts. Proceedings of the Royal Society B 278:1633–1638.

Kamal S, Grodzińska-Jurczak M, Brown G. 2015. Conservation on private land: a review of global strategies with a proposed classification system. Journal of Environmental Planning and Management 58:576–597.

Kroner REG, et al. 2019. The uncertain future of protected lands and waters. Science 364:881–886.

Laffoley D, Dudley N, Jonas H, MacKinnon D, MacKinnon K, Hockings M, Woodley S. 2017. An introduction to ‘other effective area-based conservation measures’ under Aichi Target 11 of the Convention on Biological Diversity: origin, interpretation and emerging ocean issues. Aquatic Conservation 27:130–137.

Lawler JJ, Rinnan DS, Michalak JL, Withey JC, Randels CR, Possingham HP. 2020. Planning for climate change through additions to a national protected area network: implications for cost and configuration. Philosophical Transactions of the Royal Society B 375:20190117.

McKinnon MC, et al. 2016. What are the effects of nature conservation on human well-being? A systematic map of empirical evidence from developing countries. Environmental Evidence 5:8.

Nolte C. 2020. High-resolution land value maps reveal underestimation of conservation costs in the United States. Proceedings of the National Academy of Sciences USA 117:29577–29583.

OECD (Organisation for Economic Co-operation and Development). 2019. The Post-2020 Biodiversity Framework: targets, indicators and measurability implications at global and national level. Interim Report, November version. OECD, Montreal, Canada.

Rodrigues ASL, et al. 2004. Global gap analysis: priority regions for expanding the global protected-area network. BioScience 54:1092–1100.

Rohr JR, Bernhardt ES, Cadotte MW, Clements WH. 2018. The ecology and economics of restoration: when, what, where, and how to restore ecosystems. Ecology and Society 23:15.

Sayre R, et al. 2020. An assessment of the representation of ecosystems in global protected areas using new maps of World Climate Regions and World Ecosystems. Global Ecology and Conservation 21:e00860.

Stralberg D, Carroll C, Nielsen SE. 2020. Toward a climate-informed North American protected areas network: incorporating climate-change refugia and corridors in conservation planning. Conservation Letters 13:e12712.

Strassburg BBN, et al. 2020. Global priority areas for ecosystem restoration. Nature 586:724–729.

TNC (The Nature Conservancy). 2018. Resilient and Connected Network. TNC, Washington, DC. Available from http://maps.tnc.org/resilientland/ (accessed December 2020).

UNEP-WCMC (UN Environment Programme World Conservation Monitoring Centre). 2020. World Database of Protected Areas. Protected area profile for United States of America.

UNEP-WCMC, Gland, Switzerland. Available from https://www.protectedplanet.net/country/USA (accessed December 2020).

USGS (United States Geological Survey) GAP (Gap Analysis Project). 2020. Protected Areas Database of the United States (PAD-US) 2.1. US Geological Survey data release. USGS, Washington, DC. Available from https://doi.org/10.5066/P92QM3NT (accessed December 2020).

Venter O, Magrach A, Outram N, Klein CJ, Possingham HP, Di Marco M, Watson JEM. 2018. Bias in protected-area location and its effects on long-term aspirations of biodiversity conventions. Conservation Biology 32:127–134.

Withey JC, et al. 2012. Maximising return on conservation investment in the conterminous USA. Ecology Letters 15:1249–1256.

WWF (World Wide Fund for Nature). 2020. Endorsers and Supporters. Leaders’ Pledge For Nature, Gland, Switzerland. Available from https://www.leaderspledgefornature.org (accessed February 2021).

Yang L, et al. 2018. A new generation of the United States National Land Cover Database: requirements, research priorities, design, and implementation strategies. ISPRS Journal of Photogrammetry and Remote Sensing 146:108–123.

